# Chromatin Accessibility Reveals Potential Prognostic Value of the Peak Set Associated with Smoking History in Patients with Lung Adenocarcinoma

**DOI:** 10.1101/2020.07.08.192914

**Authors:** Han Liang, Jianlian Deng, Tian Luo, Huijuan Luo, Xuan Wu, Yiwang Ye, Shubin Wang, Fuqiang Li, Kui Wu, Cong Lin

**Affiliations:** BGI-Shenzhen, Shenzhen 518083, China; China National Gene Bank, BGI-Shenzhen, Shenzhen 518120, China; Peking University Shenzhen Hospital, Shenzhen 518036, China; Guangdong Enterprise Key Laboratory of Human Disease Genomics, Shenzhen Key Laboratory of Genomics, BGI-Shenzhen, Shenzhen 518120, China; School of Future Technology, University of Chinese Academy of Sciences, Beijing, China

**Author notes:** These authors contributed equally (Han Liang, Jianlian Deng).

**Keywords:** ATAC-seq, peak, network, LUAD, smoking, prognostic, pathway

## Abstract

Considerable differences in molecular characteristics have been defined between non-smoker and smokers in patients with lung adenocarcinoma (LUAD), yet study of open chromatin patterns associated with LUAD progression caused by smoking is still lacking. Here, we constructed a novel network based on correlations between each ATAC-seq peak from TCGA data using our previously developed algorithm. Subsequently, principal component analysis was performed on LUAD samples with retained peaks filtered by the correlation network and pathway analysis was conducted for potential pathways identification. Results were verified in an independent dataset from primary LUAD samples. We identified a set of peaks that clearly differentiated long-term from short-term smokers in LUAD patients and also significantly associated with overall survival of these patients. We then investigated the gene set related to those peaks and found that the comprising genes are strongly associated with LUAD development, such as *B3GNT3, ACTN4* and *CLDN3*. They are consistent with the important roles for the associated pathways in LUAD oncogenesis induced by smoking, including glycosphingolipid biosynthesis and tight junction pathways.

In summary, our study may provide valuable insights on exploration of ATAC-seq peaks and on smoking-related LUAD carcinogenesis from a perspective of open chromatin changes.

## Introduction

Lung cancer remains the leading cause of cancer death with over 1.6 million deaths annually, and the incidence of lung cancer is still increasing worldwide (Herbst et al., 2008). More than 85% of lung cancer cases are diagnosed as non-small-cell lung cancer (NSCLC), with lung adenocarcinoma (LUAD) and lung squamous cell carcinoma (LUSC) being the two main histological subtypes. LUAD alone accounts for approximately 40% of NSCLC cases, resulting in over 500,000 deaths per year globally (Herbst et al., 2008; Herbst et al., 2018). The most important risk factor for lung cancer is still cigarette smoking, which is responsible for about 85-90% of all cases (Freedman et al., 2008; Herbst et al., 2018). The other emerging risk factors include second-hand smoking and air pollution (Oberg et al., 2011), such as PM2.5, which is claimed to cause lung cancer in many developing countries (Khilnani and Tiwari, 2018; Guo et al., 2019). Although NSCLC is strongly associated with smoking, LUAD is more common in never-smokers (Sun et al., 2007; Herbst et al., 2018). Compelling evidence indicates that never-smoker patients with LUAD have a significantly better survival rate than smokers, suggesting distinct molecular mechanisms underlying their clinical difference (Bryant and Cerfolio, 2007; Casal-Mourino et al., 2019; Lofling et al., 2019).

Recent efforts have been put to characterize different molecular alterations in LUAD using high-throughput genome sequencing, which led to a comprehensive profiling of different oncogenic driver mutations (Weir et al., 2007; Cancer Genome Atlas Research, 2014). Besides *EGFR* mutations and *ALK* fusions, for which targeted therapies have become standard treatment for LUAD, several other activated oncogenes such as, *KARS, TP53, ERBB2* and *BRAF* are also found in LUAD (Imielinski et al., 2012; Wu et al., 2015). As more and more in-depth multi-omics studies are progressing, striking differences in molecular characteristics have been discovered between LUAD arising in never-smokers and smokers. For example, LUAD patients with different levels of tobacco consumption show different mutation frequencies of the *EGFR, TP53* and *KRAS* genes, with *EGFR* mutations occurring more frequently in never smokers (Le Calvez et al., 2005; Sun et al., 2007). In addition, gene expression analysis identified distinct patterns of dysregulated genes in smokers of LUAD, of which associated altered pathways are particularly involved in the cellular immune response and cell cycle regulation(Landi et al., 2008; Liu et al., 2018). Moreover, epigenetic studies also demonstrated clear differences between methylation profiles of LUAD in never smokers and smokers(Divine et al., 2005; Toyooka et al., 2006; Alexandrov et al., 2016). To date, however, studies of open chromatin patterns associated with LUAD progression caused by smoking are still lacking.

Recently, assay for transposase accessible chromatin sequencing (ATAC-seq) has emerged as a powerful tool for profiling chromatin accessibility in different human diseases and has exerted a profound impact on understanding the coordination in gene expression processes(Buenrostro et al., 2013; Liu et al., 2019). Until now, a few studies have explored open chromatin states in NSCLC with ATAC-seq. An elegant work by Corces et al. studied chromatin accessibility of 410 tumor samples from The Cancer Genome Atlas (TCGA), including 38 cases of NSCLC(Corces et al., 2018). More recently, an integrative analysis which linked the open chromatin variations to genomic alterations among NSCLC patients provided a comprehensive open chromatin landscape of NSCLC(Wang et al., 2019). However, emphasis has not yet been placed on linking the clinical variables, such as cigarette smoking to open chromatin patterns in LUAD. In this study, we first generated a network based on correlations between peaks from ATAC-seq data of TCGA. Using retained peaks filtered by the correlation network, we then studied differences between LUAD patients with short and long-term smoking history and further identified a set of genes and their related pathways that associated with patients’ overall survival.

## Materials and Methods

### Patients and Clinical Information

Patients initially diagnosed with lung cancer at Peking University Shenzhen Hospital from July 2018 to May 2019 were included in the study. Patients’ information, including pathology, tumor type and smoking history was collected for all 6 patients (Supplementary Table 1). This study was carried out in accordance with Declaration of Helsinki and approved by the Ethics Committee of Peking University Shenzhen Hospital (Approval number: 2018033).

### ATAC-seq Library Preparation and Sequencing

Isolation of nuclei from frozen tissues and transposition reaction for ATAC-seq were performed as previously described (Corces et al., 2017). Briefly, frozen tissue sections of each sample were prepared on dry ice to avoiding thawing. Then the tissue was transferred into a pre-chilled 2-ml Dounce homogenizer containing 2 ml of 1× cold homogenization buffer. Tissues were then homogenized for several minutes with different sizes of the pestles and filtered by nylon mesh filter. The nuclei obtained by sucrose gradient centrifugation were transferred to a tube containing resuspension buffer and then pelleted by centrifugation at 500 rcf for 10 min. The supernatant was discarded and nuclei were resuspended in the Omni-ATAC-seq mix (Corces et al., 2017) containing Tn5 transposase. The reaction was incubated at 37 °C for 30 min in a thermomixer with shaking at 500 rpm and the tagmentated DNA was isolated with MinElute PCR Purification kit (Qiagen). PCR was performed using NEBNext High-Fidelity 2X PCR master mix (72°C 5min, 98°C 30sec,13cycles of: [98°C 10sec, 63°C 30sec, 72°C 1min], 72°C 5min). Sizes of PCR products were selected using Ampure XP by 0.5X volume followed by a 1.0X final volume. The libraries were then prepared based on a BGISEQ-500 sequencing platform(Huang et al., 2017) and sequenced with paired-end 50-bp read lengths.

### ATAC-seq Data Processing and Peak Calling

ATAC-seq data was processed using the ENCODE ATAC-seq pipeline^1^. Briefly, the fastq files were processed for adaptor trimming by cutadapt software (version 1.9.1)(Martin, 2011) with “cutadapt -a TCGTCGGCAGCGTCAGATGTGTATAAGAGACAG -A GTCTCGTGGGCTCGGAGATGTGTATAAGAGACAG -m 5 -O 5 -e 0.1” options, coming with the clean fastq files. Clean fastq files were aligned to the human hg38(GRCh38) genome using bowtie2(version 2.2.6)(Langmead and Salzberg, 2012) with “bowtie2 --multimapping 4 -X2000 --mm --local --nth 4” options and raw bam files were generated. Then, Samtools (Li et al., 2009) was used to sort and retain properly paired reads with “samtools -F 1804 -f 2” options; after which Picard was used to remove PCR duplicates with default arguments; mitochondrial reads were also filtered. Finally, final bam files were generated for the subsequent downstream analysis.

According to Data quality control standards of ENCODE ^2^, only samples with a transcription start site (TSS) enrichment value >5 were classified as qualified and used for downstream analysis. Peak calling was carried out to generate high quality fixed-width peaks from the final bam files in six samples, all these six samples were coupled with biological replicates to improve sensitivity and the accuracy. We called peaks using MACS2 with “macs2 callpeak -t bamfile --cap-num-peak 300000 --shift -75 --extsize 150 --nomodel --call-summits --nolambda --keep-dup all -p 0.01” options. Fixed peak set was generated by extending the peak summits position by 250 bp on either side to a final width of 501 bp, we filtered the ENCODE hg38(GRCh38) blacklist^3^ manually to avoid the interference of the high signal regions such as centromeres, telomeres, and satellite repeats, peaks that beyond the ends of chromosomes after extending were also filtered (Corces et al., 2018).

### ATAC-seq Data Analysis

We used previously described analysis method on peaks selecting and exploring(Liang et al., 2020). Briefly, we selected the available data from TCGA according to peak’s quality. A peak would be considered low-quality if it has a same value in more than 5% patients from single type of cancer, as the repeat values were likely produced by nonsense 0s before normalization. Eventually, we obtained 64316 peaks across 386 samples from TCGA dataset. To further reduce the scale of data, we then applied the previously developed algorithm on the correlation network construction with retained peaks from TCGA, in which two peaks would be connected if their direct or indirect correlation is significant(Liang et al., 2020) (Figure 1). We consider the direct correlation between two peaks is significant, if the absolute value of correlation coefficient is not less than 0.4, considering the noisy level is around 0.2. Furthermore, we consider the indirect correlation between the two peaks is significant, if their direct correlations with the third peak are both significant. Here we allow the indirect correlation to amplify peaks’ aggregation effect in the network. To assess the correlation between peaks more reliably, outlier values were removed before calculating correlation (Hearst, 1999). In practice, we used the function “cor” from R package “stats” V3.6.2 with default arguments. We selected the 10% most frequently-connected peaks for the further analysis, as those peaks were more likely to be the hub peaks in the network with high connectivity.

**Figure 1.**
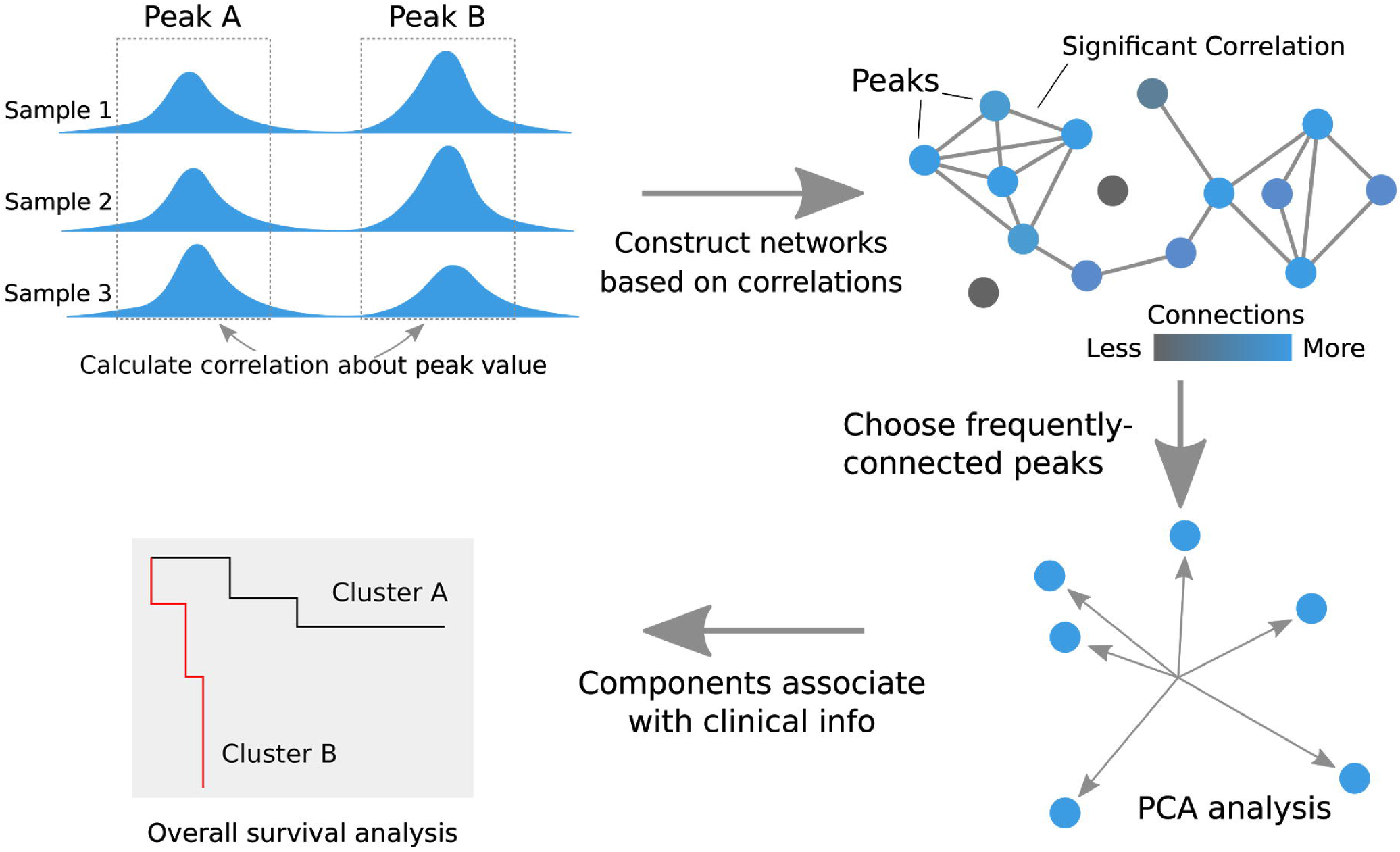
The graphical abstract of analyses performed in this study. To reduce the complexity of the large data and only emphasize on important peaks, a correlation network was constructed. Peaks were connected to each other if their direct or indirect correlations are significant (blue dots). We chose the 10% most frequently-connected peaks as the important peaks for the further PCA analysis. We then analyzed the association between components obtained from PCA and patient’s overall survival.

The unsupervised classification method principal component analysis (PCA) was used to analyze the selected ATAC-seq peaks from 22 LUAD patients in TCGA dataset. We used the function “PCA” from R package “FactoMineR” V1.34 with default arguments which produced 5 most important components(Lê et al., 2008). The association between components and smoking status were checked by the distribution of samples classified by components. To statistically assess the difference between distribution of samples from LUAD patients with different smoking histories (group “1-2 Years” and the group “3-4 Years”), we used the Wilcox Test. We used the function “survdiff” from R package “survival”(Harrington and Fleming, 1982), in which Chi-Squared Test was used to assess the overall survival analysis. Then the function “pchisq” from R package “stats” was used to calculate P value with arguments df=1, lower tail=F.

### Pathway Analysis

For the pathway analysis, all linc-RNA genes (such as RP11-genes) have been removed as they are excluded in KEGG database. We searched the potential pathways using DAVID 6.8. with the non-linc-RNA genes related to top 100 highest contributing peaks of targeted component. Three pathways were found with p value less than 0.1, which is the recommended value of DAVID (Huang da et al., 2009).

## Results

### A correlation network is built based on ATAC-seq data from TCGA

High-quality ATAC-seq data of 410 tumor samples across 23 cancer types was downloaded and collected from TCGA. The extended peak summits with fixed width of 501 bp were extracted from the dataset and used to perform the further analysis(Corces et al., 2018). We first constructed a correlation network with retained peak summits from TCGA, in which two peaks would be connected if their peak values were correlated with each other significantly (Figure 1). To this end, correlations between each peak value in all samples were calculated and the 10% most frequently-connected peaks were chosen after removing the low-quality peaks and outliers. Using PCA on those chosen peaks across all cancer types, we identified distinct clusters labeled based on the different cancer-type enrichment (Figure 2). We found strong concordance between our clustering results and the t-distributed stochastic neighbor embedding (t-SNE) results from TCGA (Corces et al., 2018).

**Figure 2.**
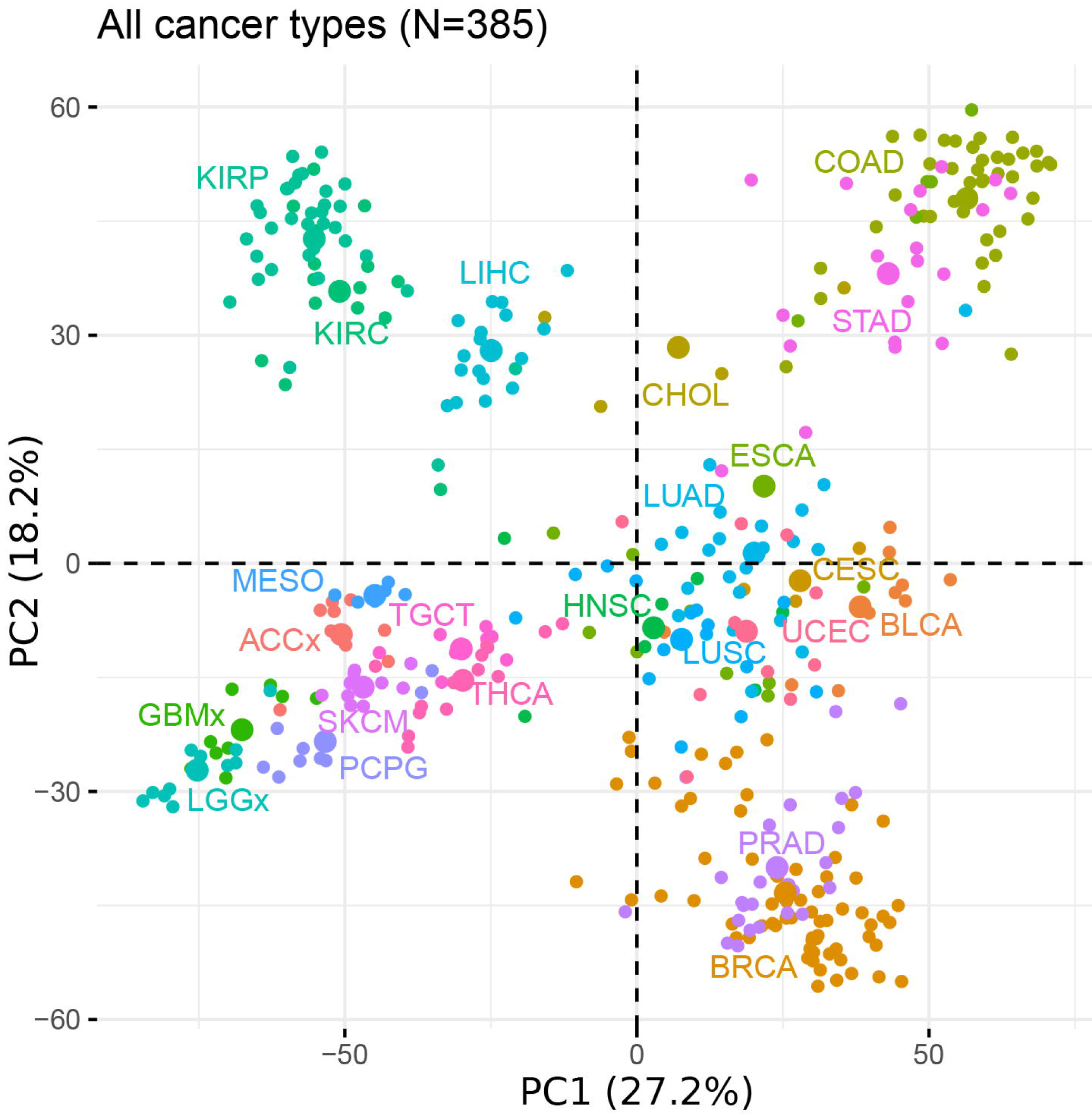
Principal Component Analysis on the selected peaks from TCGA ATCA-data. The unsupervised PCA for the 10% most frequently-connected peaks selected from TCGA ATAC-seq data across all cancer types. Each dot represents a given sample. Color represents the cancer type shown in the Figure

### Identification of the smoking associated peak set in LUAD patients and related pathways

We next focused on the analysis of 22 LUAD patients in the TCGA dataset, using the same correlation network and analyzing methods described above. PCA result indicated that the first principal component (PC1) explained 16.9% of the variability, while PC2 explained 12.8% of the variability in the peaks with all LUAD samples (Figure 3A). The LUAD samples do not appear to form distinct patterns within the two dimensions generated by PCA analysis, interestingly however, samples with a shorter smoking history seemed to be closer to PC2 axis (Figure 3A). A sample has a shorter distance towards PC2 axis means its PC2-related peaks are less variable. Using each sample’s absolute distance from the PC2 axis, we compared the difference between the shorter and longer smoking history groups (1-2 years vs. 3-4 years) (Figure 3B). The results showed that the group with 3-4 yeas of smoking indeed has a significant longer absolute distance from PC2 axis compared to the group with 1-2 years of smoking, suggesting the ATAC-seq peaks associated with PC2 are influenced by smoking history. Additionally, shorter absolute distance from PC2 axis, representing more stable PC2-related peak values, is significantly associated with better overall survival, indicating a potential prognostic value for the corresponding peaks of PC2 (Figure 3C). We thus further studied the gene set related to those peaks according to the defined peak-gene relationships (Corces et al., 2018)(Supplementary Table 2) and explored the associated pathways. Consequently, we identified three potential pathways, including glycosphingolipid biosynthesis, tight junction and metabolic pathways (Table 1).

**Figure 3.**
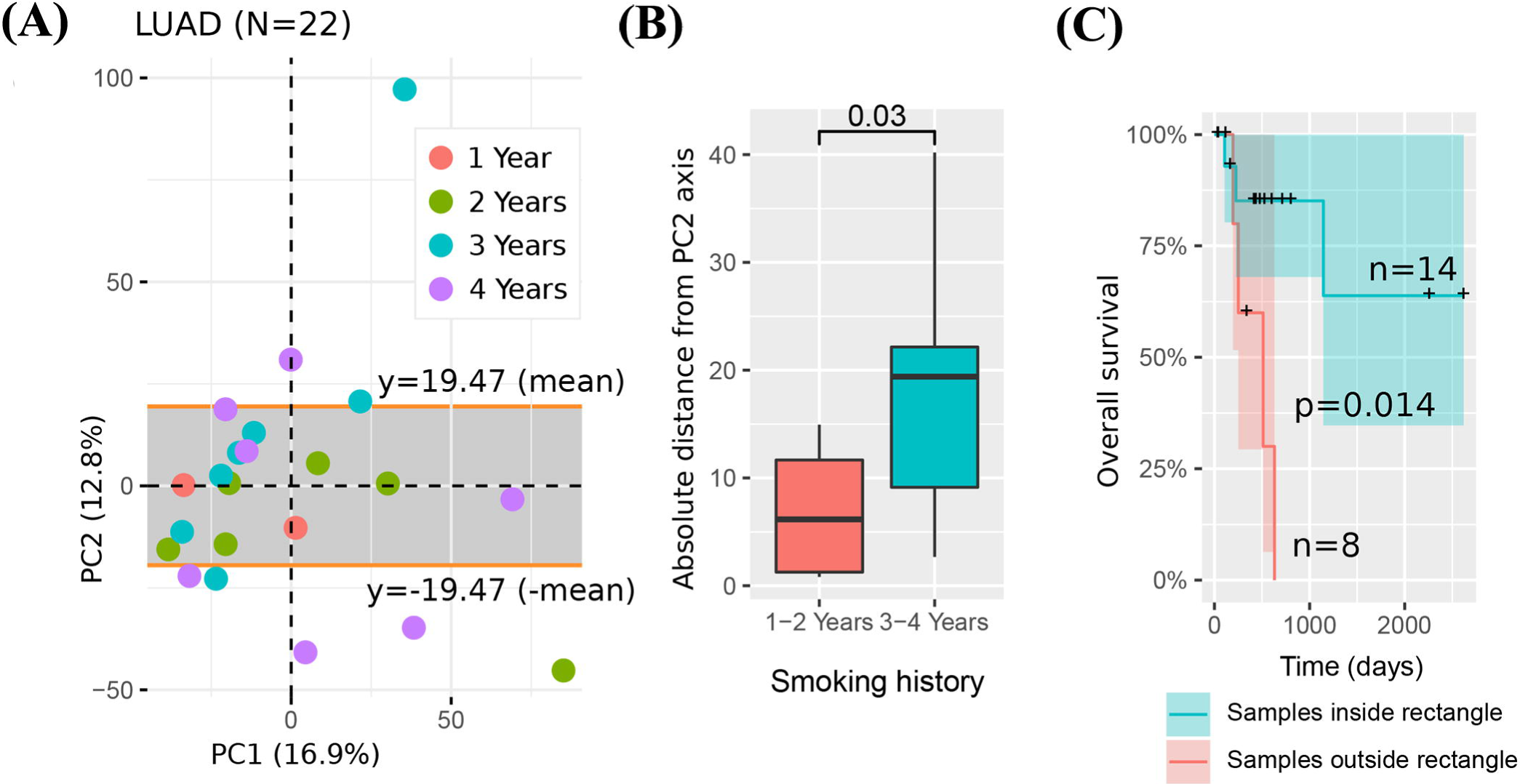
Identification of the smoking associated peak set from LUAD patients using TCGA ATCA-data. **(A)** The unsupervised PCA on LUAD samples (N=22) from TCGA. Dots present samples and its colors present patients’ different smoking histories. The upper and lower sides of the gray rectangle area are defined by the ±mean of all LUAD samples’ distances from the x axis. **(B)** The absolute distance from PC2 axis of LUAD patients with different smoking histories. We divide samples into groups of 1-2 years (N=7) and 3-4 years (N=13). Two outliers, one from each group have been removed respectively. Each sample’s absolute distance from PC2 axis (the line y=0 in Figure 3A) was measured and Wilcox test was performed to compare the difference between the two groups. The difference is statistically significant as P < 0.05. **(C)** The overall survival (OS) difference between LUAD patients with different absolute distance towards PC2. Samples were divided into two groups based on whether it locates in the gray rectangle in Figure 3A, and the OS difference between the group with samples outside rectangle (N=8) and the group with samples inside the rectangle (N=14) was measured by Chi-Squared Test. The difference is statistically significant as p value=0.014. Patients survived but stopped being tracked are indicated by *crosses*.

**Table 1.** Three pathways related to PC2 high contributing peaks found with DAVID 6.8.

| Pathways | p value | Genes |
| --- | --- | --- |
| hsa00601: Glycosphingolipid biosynthesis - lacto and neolacto series | 0.0080 | *B4GAT1, FUT3, B3GNT3* |
| hsa04530: Tight junction | 0.010 | *ACTN4, CLDN3, TJP3, LLGL2* |
| hsa01100: Metabolic pathways | 0.091 | *DGKA, GALNT3, POLE2, HSD17B2, B4GAT1, AKR1B1, FUT3, B3GNT3, GALE, AGMAT, GAL3ST1* |

### Verification of the smoking associated peak set in primary LUAD samples

To verify the smoking associated peak set we identified, we collected primary tumors from 6 patients with LUAD and performed ATAC-seq for each sample. Each of the ATAC-seq data generated 48∼246M (median = 108M) reads. We then filtered out 2 samples which did not reach TSS value of 5, leaving 4 samples from LUAD patients for the following analysis (Supplementary Table 3). We calculated their PC2 values according to weights of PC2-related peaks showed in LUAD samples from TCGA data, so that the PC2 values of these samples resembled that seen as absolute distance from PC2 axis in Figure 3A. The result show that the one sample with smoking history has higher absolute PC2 values compared to the other three smoking-free samples (Figure 4), which is in agreement with our previous observation from TCGA ATAC-seq data.

**Figure 4.**
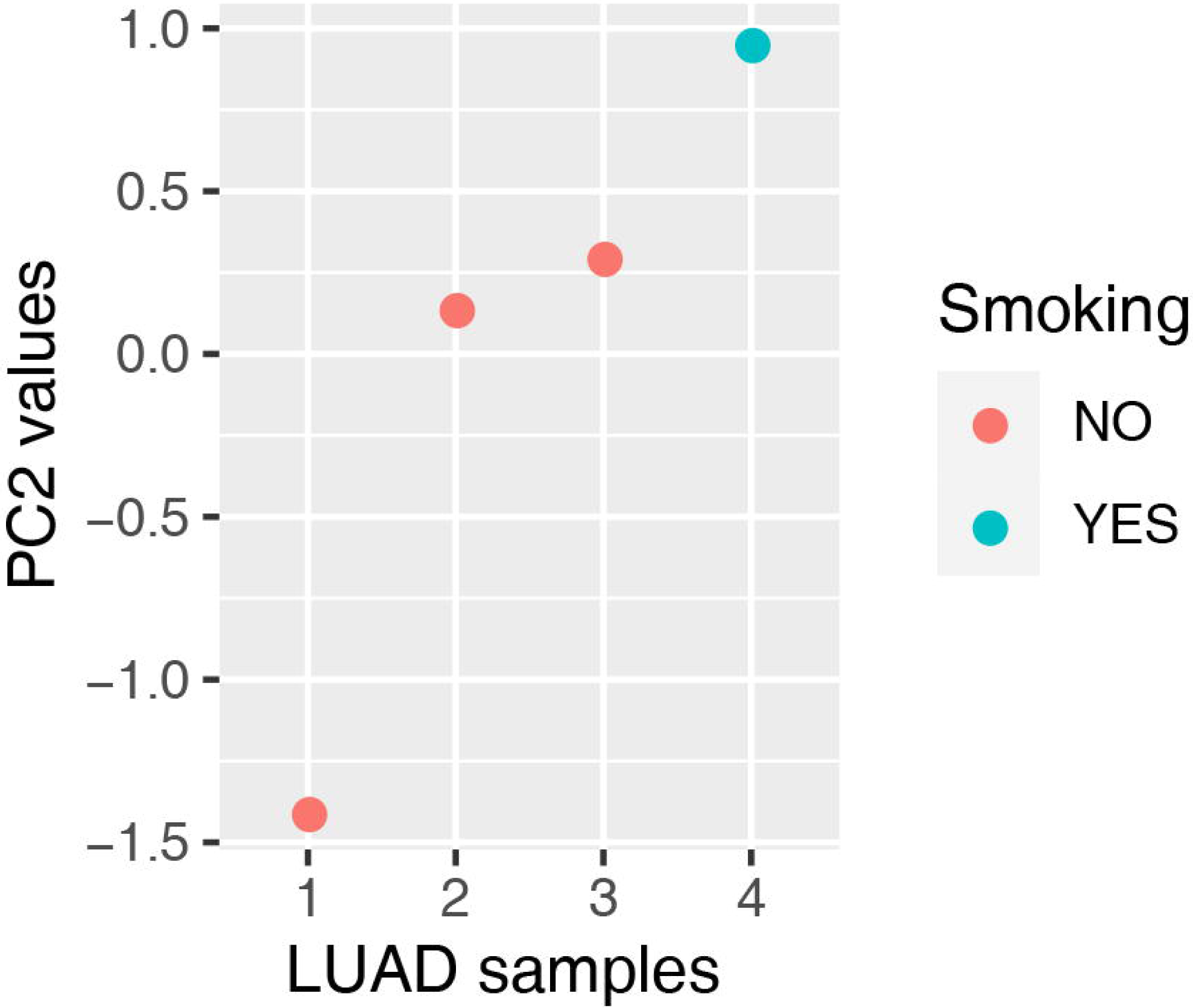
Verification of the smoking associated peak set in primary tumors collected from patients with LUAD. We performed analysis on 4 LUAD patients, one of which has smoking history (blue dot), whereas other three are non-smokers (red dots). Their PC2 values were calculated by weights of peaks associated with PC2 axis in the LUAD samples from TCGA. The PC2 values were normalized into a suitable range and resembled that seen as absolute distance from PC2 in Figure 3A.

## Discussion

During the past decade, many important differences in genomes and signaling pathways between non-smokers and smokers with LUAD were unveiled by in-depth analyses, yet the precise chromatin accessibility alterations induced by smoking remained obscure(Toyooka et al., 2006; Sun et al., 2007; Landi et al., 2008). In the present study, we first constructed a correlation network with ATAC-seq data from TCGA using the algorithm we previously developed, and analyzed 22 LUAD samples with peaks selected by the network. We determined smoking related peaks with potential prognostic value and found associated pathways based on the defined peak-gene relationships.

Different from the classic analysis on correlation of gene expression and chromatin accessibility (Corces et al., 2018; Wang et al., 2019), we solely focused on correlations between each peak and constructed the network based on the plausible theory that peaks which are highly connected with many other peaks would be more likely to have a crucial function in gene regulatory processes. The correlation network assisted us with critical peaks selection and enabled a more precise analysis on further peak classification. With this novel conception, smoking-related peak set in this study was effectively identified from the massive ATAC-seq data. The effect of this method was also reflected by the identification of specific mitosis-related expression pattern in the previous work on data mining of transcriptomes from 5,001 cancer patients cross 22 cancer types(Liang et al., 2020). Therefore, we believe that this method is practical in revealing crucial factors from complicated datasets and will additionally increase analytical effects on studies of future multi-omics data.

The peaks selected by the correlation network were further analyzed with PCA and we demonstrated that the variability of PC2-related peaks obtained from LUAD patients was influenced by smoking history. Intriguingly, PC2 was also associated with overall survival of these LUAD patients (Figure 3), pointing to genes that corresponding to PC2-related peaks as potential effectors in carcinogenesis of LUAD. Indeed, the pathways significantly associated with PC2-related gene set, including glycosphingolipid biosynthesis and tight junction pathways, are proven to be important in tumor progression(Soini, 2012; Zhuo et al., 2018). Glycosphingolipid is a crucial part of the cell membrane, which is formed by a ceramide backbone linked to a glycan moiety. The aberrations in glycosphingolipid metabolism can lead to abnormal cell rearrangements and thus contribute to proliferation, metastatic cellular invasion and even resistance to chemotherapy in tumor cells (Ogretmen, 2018; Russo et al., 2018; Zhuo et al., 2018). In the context of LUAD progression, several previous findings showed that the exposure to cigarette smoking results in increased levels of ceramide, which in turn promotes aberrant EGFR activation and leads to elevated tumorigenic signaling(Goldkorn et al., 2013). Additionally, dysregulated expressions of genes that are related to glycosphingolipid pathways, for instance *B4GAT1* and *B3GNT3* identified by PC2-related peaks, have also been shown to correlated with the observed malignancy(Willer et al., 2014; Gao et al., 2018). Among those genes, overexpression of *B3GNT3* is specifically associated with unfavorable overall survival in NSCLC patients(Gao et al., 2018). As for the tight junction pathway, alterations in expressions of its proteins induced by smoking are involved in increased lung epithelial permeability and epithelial-mesenchymal transition (EMT), thus influencing cancer progression(Shaykhiev et al., 2011; Soini, 2012). The genes we identified through PC2-related peaks, such as *ACTN4* and *CLDN3*, are consistent with the important role for this pathway in LUAD oncogenesis(Zhang et al., 2017; Tentler et al., 2019). Specifically, *ACTN4* amplification can be induced by smoking and is also associated with increased mortality risk among patients with early stage of LUAD(Noro et al., 2013). Therefore, the gene set we identified through ATAC-seq peaks comprises genes that are strongly associated with smoking status and may have potential prognostic value in patients with LUAD. However, before drawing conclusions on potential clinical implications of these genes, it is important to elucidate their function with respect to LUAD development.

The PC2-realted peaks we identified from TCGA ATAC-seq data effectively differentiated long-term from short-term smokers in LUAD tumors. To confirm this observed correlation between the peak-set and smoking status, we tried to restored the values of PC2-related peaks in an independent dataset from primary LUAD samples. Although the smoker was clearly distinguished from the other three non-smokers in the LUAD samples by the PC2-related peaks, due to the small sample size, the results cannot reach statistical significance. Further studies are needed with datasets of LUAD patients to confirm the effects of PC2-related peaks exerted in this study.

In conclusion, our study provides a novel method to explore ATAC-seq peaks and identifies a set of peaks with potential prognostic value based on chromatin accessibility alterations induced by smoking in LUAD patients. These findings provide information on smoking-related LUAD carcinogenesis from a perspective of open chromatin changes and may have impact on future clinical application, although further assays are warranted to confirm these findings.

## Data Availability

The data that support the findings of this study have been deposited into CNGB Sequence Archive (CNSA: https://db.cngb.org/cnsa/) of CNGBdb with accession number CNP0001103.

## Author Contributions

CL, HL, and JD conceived the study. HL and JD performed the ATAC-seq data analysis; TL and HJL carried out the experiments; XW, YY and SW conducted sample collection; FL contributed to the ENCODE pipeline localization; KW instructed the study; CL supervised the study and wrote the manuscript.

## Acknowledgement

This work was supported by China National GeneBank.

## Footnotes

1 https://www.github.com/ENCODE-DCC/atac-seq-pipeline

2 https://www.encodeproject.org/atac-seq/. Retrieved 24/12, 2019

3 https://www.encodeproject.org/files/ENCFF356LFX/

